# *Mid1* deletion leads to cognitive dysfunction in Opitz syndrome by regulates neural rhythms through the inhibition of p-Creb by PP2Ac

**DOI:** 10.1101/2024.10.09.617363

**Authors:** Ziye Yang, Pengxiang Li, Yue Chen, Xiaoyu Guo, Ping Liu, Guangjian Ni, Shuang Liu, Liqun Chen, Dong Ming

## Abstract

MID1 is an E3 ubiquitin ligase of the tripartite motif (TRIM) subfamily of RING-containing protein^1^. MID1 is involved in many basic biological processes, especially during embryonic development. Mutation, truncation or complete deletion of *MID1* gene is the cause of Opitz G/BBB syndrome (OS). OS is a rare genetic disease of nervous system, which is characterized by midline structural development defects during embryogenesis, including structural brain abnormalities, developmental retardation and mental retardation^2^. Although the function of *MID1* has been studied for many years, the effect and mechanism of complete deletion of *MID1* gene on OS nervous system still need to be further explored. Here we find that *Mid1* gene is necessary for the normal development of hippocampus (HPC), and *Mid1* gene knockout (*Mid1*^*-/y*^) mice showed a significant decrease in α rhythm in HPC and abnormal synchronization of γ rhythm in prefrontal cortex and hippocampus (PFC-HPC), showing decreased synaptic plasticity and learning and memory dysfunction.

## Results and Discussion

It has been reported that OS shows developmental and motor retardation^3^. In this study, the body weight and the weight of various organs of *Mid1*^*-/y*^ were tested, and it was found that the average body weight of *Mid1*^*-/y*^ were lower than that of *Mid1*^*+/y*^ during growth, but there was no significant difference (Supplementary Fig. S1a), and there was no significant difference in the weight of liver, lung, kidney and brain in *Mid1*^*-/y*^ (Supplementary Fig. S1b-e). Monitoring the movement behavior of *Mid1*^*-/y*^ within 24h showed that the total movement distance, average movement speed and number of standing were decreased (Supplementary Fig. S1f-k). In order to further study the effect of *Mid1* on nervous system, we first carried out elevated plus maze, open field and tail suspension tests, and found no negative effect of *Mid1*^*-/y*^ on anxiety and depression. (Supplementary Fig. S2). However, in the Y-maze experiment, the spontaneous alternation score of *Mid1*^*-/y*^ was significantly reduced by 21.0±9.5% (Fig. 1a). In the morris water maze, it was found that after the fifth day of continuous training, the time required for *Mid1*^*-/y*^ to find the platform was significantly increased by 81.4±38.2%. (Fig. 1b; Supplementary Fig. S3a). On the sixth day of the test, it was found that the total distance, staying time and crossing times of *Mid1*^*-/y*^ in the platform area decreased (Fig. 1c, d; Supplementary Fig. S3b-e). These results show that *Mid1*^*-/y*^ will lead to the decline of learning and memory ability.

**Fig. 1:**
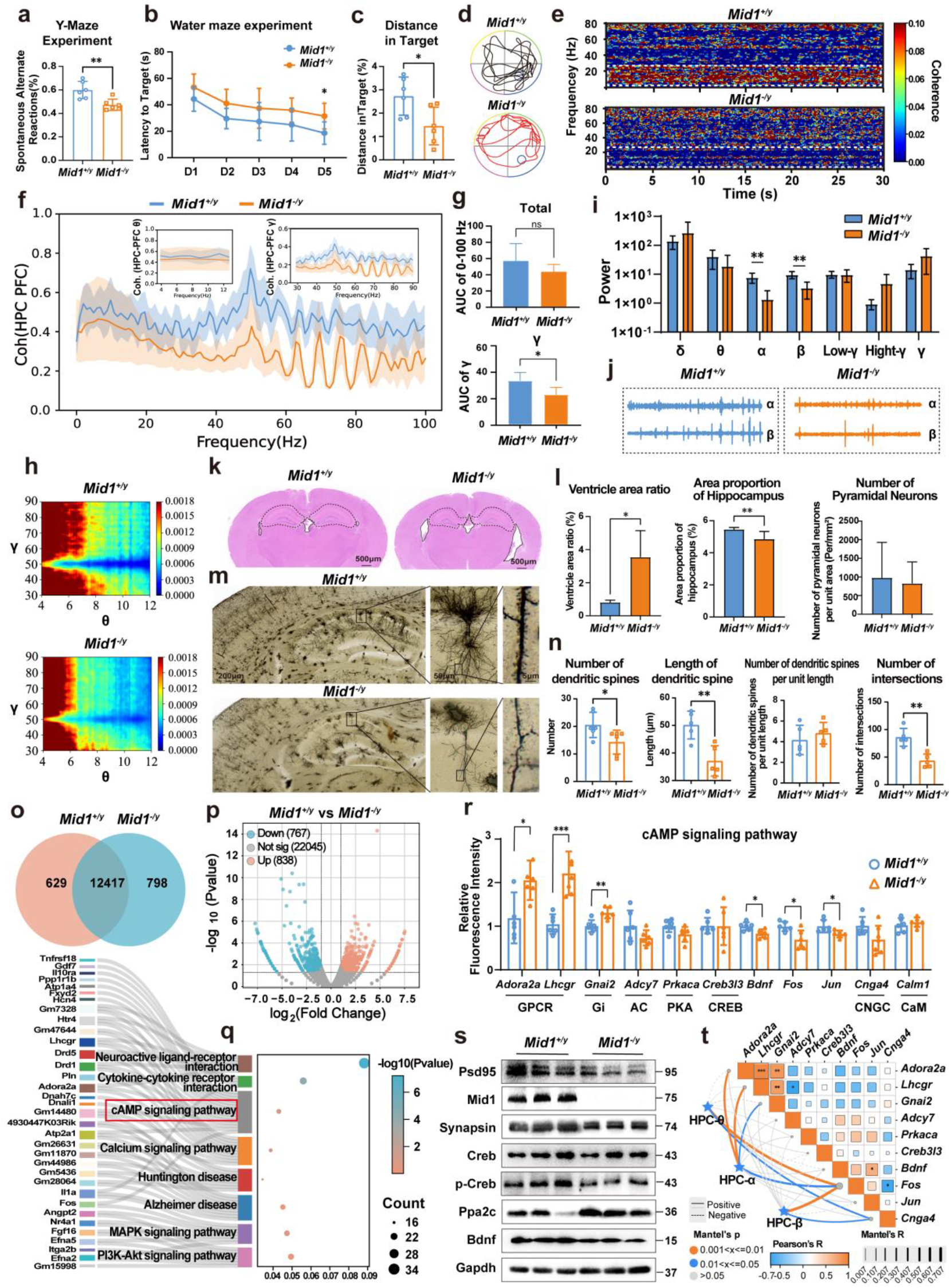
X-linked microtubule-associated protein *Mid1* regulates hippocampal development affects neural rhythm, learning and memory. **a** The spontaneous alternate scores of *Mid1*^*+/y*^ and *Mid1*^*-/y*^ in Y maze test (*n* = 6 mice). **b** The time required to find the platform by training *Mid1*^*+/y*^ and *Mid1*^*-/y*^ for five consecutive days in the water maze test (*n* = 6 mice). **c** The total distance of *Mid1*^*+/y*^ and *Mid1*^*-/y*^ moving in the platform area in water maze test (*n* = 6 mice). **d** Images of representative tracks of the *Mid1*^*+/y*^ and *Mid1*^*-/y*^ tested on the sixth day of the water maze test. **e** Time-frequency coherence between HPC and PFC in *Mid1*^*+/y*^ and *Mid1*^*-/y*^ (*n* = 6 mice). **f-g** The synchrony between the PFC and HPC in the frequency range of 0-100 Hz in *Mid1*^*+/y*^ and *Mid1*^*-/y*^ (*n* = 6 mice). **h** θ-γ cross-frequency coupling in HPC-PFC in *Mid1*^*+/y*^ and *Mid1*^*-/y*^, respectively (*n* = 6 mice). **i** Power data of individual rhythms in HPC of *Mid1*^*+/y*^ and *Mid1*^*-/y*^ under different rhythms (δ:0.5∼3.5 Hz, θ:4∼12 Hz, α:8∼13 Hz, β:14∼35 Hz, γ:30∼90 Hz, Low-γ:30∼55 Hz) (*n* = 5-6 mice). **j** The representative signals of α and β rhythm frequencies in the HPC of *Mid1*^*+/y*^ and *Mid1*^*-/y*^. **k** Representative pictures of HE staining of brain tissue of *Mid1*^*+/y*^ and *Mid1*^*-/y*^. **l** The ventricle area, HPC area and the number of HPC pyramidal neurons in *Mid1*^*+/y*^ and *Mid1*^*-/y*^ were counted (*n* = 6 mice). **f** Representative pictures of golgi staining in *Mid1*^*+/y*^ and *Mid1*^*-/y*^. **g** Changes of the number of dendritic spines, the length of dendritic spines, the number of dendritic spines per unit length and the number of intersections between dendrites and concentric circles in *Mid1*^*+/y*^ and *Mid1*^*-/y*^ (*n* = 5 mice). **o** Venn diagram of common and specific genes detected by transcriptome of *Mid1*^*+/y*^ and *Mid1*^*-/y*^ HPC. **p** Differential gene volcano map. The abscissa represents the change of gene expression multiple in *Mid1*^*+/y*^ and *Mid1*^*-/y*^, and the ordinate represents the significant level of gene expression difference between the two groups. **q** Sankey and bubble diagram reflects the main pathways enriched by KEGG, and the significant differential genes contained in each pathway. **r** The mRNA levels of gene in the HPC were tested via RT-qPCR by normalization to the level of *β-actin* (*n* = 6 mice). **s** Western Blot results of protein in HPC of *Mid1*^*+/y*^ and *Mid1*^*-/y*^ (*n* = 6 mice). **t** Pairwise comparisons of important genes in the pathway are shown, with a color gradient denoting Spearman’s correlation coefficients. Different rhythms (θ, α, β) of HPC was was related to the effect of different genes by partial Mantel tests. Edge width corresponds to the Mantel’s r statistic for the corresponding distance correlations, and edge color denotes the statistical significance based on permutations. * indicates highly significant correlation (P < 0.05), ** indicates highly significant correlation (P < 0.01), *** indicates highly significant correlation (P < 0.001).

Neuroelectrophysiological technology provdes a more detailed and in-depth method for studying learning and memory ability through its advantages of direct measurement of neural activity, high time sequence resolution and objectivity^4^. There is a strong neural synchronization between HPC and PFC to regulate various cognitive functions^5^. But we found that the time-frequency correlation between the two brain regions of *Mid1*^*-/y*^ decreased (Fig. 1e), and the coherence between HPC and PFC at 0-100 Hz frequency was also weakened, in which the γ correlation was significantly decreased by 33.2±16.0% (Fig. 1f, g). The cross-coupling degree of θ-γ rhythm in HPC and PFC related to learning and memory also decreased (Fig. 1h)^6^. We analyzed the neural rhythms in the HPC and PFC of *Mid1*^*-/y*^, respectively, to determine the important brain regions affected by *Mid1*. Through time-frequency analysis and power spectral density quantitative analysis, it was found that the HPC power of *Mid1*^*-/y*^ decreased at 0-100 Hz (Supplementary Fig. S4a, b). Among them, the power density of α and β rhythm in HPC decreased significantly by 83.3±10.9% and 57.7±30.9%, respectively (Fig. 1i), and the representative waveform diagram is shown in Fig. 1j. However, no significant changes were found in the PFC (Supplementary Fig. S4c, d). To sum up, the results of this part show that *Mid1*^*-/y*^ have decreased synchronization between HPC-PFC of 30-90 Hz, and abnormal neural rhythm in HPC of 8-35 Hz.

To further explore the physiological and pathological effects of *Mid1* on HPC, hematoxylin-eosin staining results showed that the area of ventricles in *Mid1*^*-/y*^ were significantly increased by 3.5±1.5 times and the area of HPC were significantly decreased by 15.1±6.6%, but the number of HPC pyramidal neurons did not change significantly (Fig. 1k, l). In order to determine whether the abnormality of brain tissue structure and motor behavior in *Mid1*^*-/y*^ were related to inflammation, we tested the whole body thermal imaging of *Mid1*^*-/y*^, and there was no significant change (Supplementary Fig. S5a). The analysis of whole blood showed that the contents of red blood cells and white blood cells (neutrophils, lymphocytes, eosinophils, basophils and monocytes) had no obvious change (Supplementary Fig. S5b-i). These results suggest that the abnormalities in physiological structure and motor behaviour seen in *Mid1*^*-/y*^ are not related to inflammation and may be closely related to their own development. Learning and memory are complex processes of neural activity, involving information transmission and synaptic connections. Golgi staining showed that although the number of dendritic spines per unit length in the HPF of *Mid1*^*-/y*^ did not change significantly, the total number and the length of dendritic spines decreased significantly by 30.3±18.0% and 26.8±11.8% respectively, and the intersection points of dendrites and concentric circles decreased (Fig. 1m, n). Immunofluorescence staining showed that the positive signals of synapse markers Syn and Psd95 in *Mid1*^*-/y*^ were obviously weakened, and there was no obvious change in microglia labeled with Iba1 (Supplementary Fig. S5a-c).

In order to fully understand the changes of HPC gene level in *Mid1*^*-/y*^, we carried out RNA-seq analysis. The results showed that *Mid1*^*+/y*^ and *Mid1*^*-/y*^ detected the same 12,417 genes (Fig. 1o), including 838 genes significantly up-regulated and 767 genes down-regulated (Fig. 1p). Gene annotation analysis enriched by the kyoto encyclopedia of genes and genomes (KEGG) pathway shows that the neuroactive ligand-receptor interaction, cAMP signaling pathway, calcium signaling pathway and other pathways have changed significantly (Fig. 1q). Then, we examined the changes of mRNA levels of genes in the cAMP pathway. The results showed that the mRNA levels of HPC adenosine A2a receptor (*Adora2a*), luteinizing hormone (*Lhcgr*) and G protein subunit α I2 (*Gnai2*) in *Mid1*^*-/y*^ increased significantly, while the mRNA levels of brain-derived neurotrophic factor (*Bdnf*), fos proto-oncogene (*Fos*) and jun proto-oncogene (*Jun*) decreased significantly (Fig. 1r). Many studies have proved that *Mid1* knockdown will increase the accumulation of protein of PP2Ac^2,7^. However, PP2Acα promoter defines a cAMP response element (CRE) site flanked by CpG motifs and that methylation controls the binding of p-CREB and the activity of the promoter^8^, thus regulating gene transcription and widely participating in the learning and memory process of the nervous system^9^. Therefore, we further verified at the protein level that the deletion of *Mid1* gene significantly increased the accumulation of Pp2ac protein and inhibited the activity of p-Creb, which led to the decrease of Bdnf protein expression level in the downstream cAMP pathway, and finally affected the growth, morphology and function of axons and dendrites of hippocampal neurons, resulting in the decrease of synaptic density (Fig. 1s, Supplementary Fig. S5d). In addition, we analysed the correlation between rhythms in different brain regions and behavioural alterations in *Mid1*^*-/y*^ and found that HPC θ, α, β were significantly correlated with abnormal behavioural changes (Supplementary Fig. S6). Mantel tests were performed on the genes that changes in the pathway and found that the series of alterations caused by *Mid1* gene deletion correlated most significantly with HPC α rhythm changes (Fig. 1t).

Generally speaking, our research shows that *Mid1*^*-/y*^ are characterized by a significant decrease in HPC α rhythm and a significant decrease in γ correlation betweenHPC and PFC. The main reason is that the deletion of *Mid1* gene will increase the accumulation of Pp2ac protein, inhibit the activity of p-Creb, affect the downstream cAMP pathway, lead to the decrease of synaptic density and plasticity, and ultimately affect the learning and memory ability. This is helpful to understand the causes of mental retardation in OS and is of great significance for the diagnosis and treatment of the disease.

## MATERIALS AND METHODS

*Mid1* KO mice, a kind gift from Zhiqi Xiong and Yiguo Wang, were genotyped as described previously^1^. *Mid1*^*+/y*^ and *Mid1*^*-/y*^ mouse genotyping results are showed in *SI Appendix*, Fig. S7. All the animal experiments conducted in this work gained the approval of the Animal Ethics Committee of Tianjin University (Approval No. TJUE-2024-164).

Neuroelectro physiological experiment. In prefrontal cortex and hippocampus (PFC-HPC), *in vivo* field potentials were recorded simultaneously. The mice were placed in a stereotactic frame and treated with isoflurane. When the 1 cm long skin incision was generated, each soft tissue from the skull surface was removed; and an electric cranial drill was used to drill the two small holes with a diameter of 1 mm. Two sections (each with four tungsten wires closely spaced) of tungsten recording electrodes were placed in the PFC (1.2 mm mediolateral and 2.2 mm anterior to Bregma, depth from the dura, 1.4 mm) and HPC (0.4 mm mediolateral and 1.9 mm posterior to Bregma, depth from the dura, 2.5 mm) brain regions. In the PFC-HPC regions, the LFP signals were collected for 5 min simultaneously. In this paper, the LFP data used were collected by using (Blackrock, USA) a sampling frequency of 20 kHZ.

See the Supplementary Information for a full description of Materials and Methods.

## ACKNOWLEDGEMENTS

We thank all the members in our Lab for their great assistance with this study. This work was supported by the National Key Research and Development Program of China (grant numbers: 2022YFF1202900), the National Natural Science Foundation of China (grant numbers: 82471196). We thank Zhiqi Xiong from the Chinese Academy of Sciences and Yiguo Wang from Tsinghua University for providing us with *Mid1* KO mice.

## AUTHOR CONTRIBUTIONS

L.C. C.Y. initiated the project, conceived and supervised the research; Z.Y., X.G. and P.L. performed the experiments; P.L. and Z.Y. analyzed the data; L.C. and Z.Y. wrote the manuscript with input from all authors.

## DATA AVAILABILITY

The source data underlying the respective main text (Figs. 1) and Supplementary Figures are provided as Source Data Files. Source data are provided in this paper. The source data for Fig. 5 were deposited on the web server http://www.ncbi.nlm.nih.gov/bioproject/1131272. Please use the BioProject ID PRJNA1131272.

## CONFLICT OF INTEREST

The authors declare no competing interests.

**Correspondence** and requests for materials should be addressed to Liqun Chen.

